# Environment and disease have tissue-specific effects on the tree microbiome

**DOI:** 10.1101/2025.08.15.670310

**Authors:** Jim Downie, Alejandra Ordonez, Marine C. Cambon, Usman Hussain, Nathan Brown, Manfred Beckmann, Jasen Finch, John Draper, Sandra Denman, James E. McDonald

**Author notes:** Corresponding author: James McDonald.

## Abstract

Trees are essential for ecosystem function, but due to their long lifespan, are disproportionately impacted by climate change and disease. Tree-associated microbiota are critical for tree health and resilience, but the composition and function of tree microbiomes across different tissue types, and how environmental factors and disease impact tree microbiomes, is poorly understood. Oak trees are major constituents of forests of the Northern hemisphere, but are increasingly impacted by climate and disease. Here, we studied the oak microbiome across Britain, combining 16S rRNA gene and ITS microbial community profiling and shotgun metagenomics of leaf, stem and root/rhizosphere samples, and developed a three-level occupancy model to describe microbiota distribution across the landscape. We show that oak leaf, stem and root/rhizosphere tissues harbour taxonomically and functionally distinct microbiota and identified differences in tissue-specific effects of environmental variables (e.g. temperature, rainfall, ion deposition) on microbiome composition and function. We generated 1657 bacterial, archaeal and fungal metagenome-assembled genomes representing key members of the oak microbiome. Furthermore, the stem microbiome of oak trees with symptoms of Acute Oak Decline, a complex decline disease driven by abiotic and biotic stressors, exhibited reduced bacterial and fungal richness and altered microbiome function. This work represents the most comprehensive microbiome study of a tree species to date. Understanding how tree-associated microbiota respond to environmental change and disease across different tissues is crucial to predict future climate and disease impacts on tree microbiome function, and inform translational approaches to modulate tree microbiomes for plant health.

## Introduction

Trees represent the largest component of living biomass on Earth ^1^ and provide essential ecosystem, societal and economic services that sustain planetary health and shape the terrestrial biosphere ^2^, including CO_2_ assimilation ^3^ and atmospheric methane uptake ^4^. However, trees face increasing threats from environmental change anthropogenic activities, and the emergence of novel pests and diseases ^5,6^. Due to their sessile nature and long lifespan, trees may have a reduced capacity to adapt to these changes, or to coevolve with encountered pathogens ^7^. By contrast, plant-associated microorganisms can respond over much shorter time scales, due to their short generation time and potential for wide dispersal. These micro-organisms have been shown to play critical roles in tree health through nutrient acquisition, growth promotion, inducing stress tolerance, and protection from disease. Consequently, plant-associated microbiota could play especially critical roles in host adaptation to stressors, extending phenotypic adaptability in the host ^8,9^.

Tree-associated microbiota extend the host phenotype via numerous mechanisms ^6^; root-associated mycorrhizal fungi enhance nutrient acquisition by the host tree ^10^ and rhizosphere and endosphere bacteria stimulate plant development and growth via the production of phytohormones such as auxins, gibberelins, and ethylene ^11^. In crop plants, inoculating maize with strains of *Burkholderia* and *Enterobacter* increased plant biomass, photosynthetic efficiency, and improved tolerance to drought stress through enhancing water retention ^12^, demonstrating the beneficial properties of plant-associated microbiota. As global tree species experience escalating outbreaks of disease by pests and pathogens, microbial-mediated protection from disease, either through direct interactions, or indirectly via competition or modulation of host responses, is increasingly important. A strain of *Luteimonas* bacteria found on healthy ash (*Fraxineus excelsior*) trees suppressed the growth of *Hymenoscyphus fraxineus* (the causal agent of ash dieback) when inoculated onto seedlings, and resulted in reduced disease symptoms that lasted over a year ^13^. The composition and function of microbial communities in tree tissues is therefore a major determinant of tree health, but arguably, the majority of research has focussed on the rhizosphere, and the impact of environmental variables and disease on the composition and function of the tree microbiome across different tissue types is poorly understood, limiting our understanding of the impact of climate stressors on tree health and hampering opportunities to develop tools to modulate tree microbiomes to promote health and suppress disease.

Oak trees (*Quercus spp.*) are major constituents of temperate and tropical forests of the Northern hemisphere ^14^, and are ecologically, economically and culturally important, providing essential roles in supporting biodiversity, sequestering carbon and climate regulation ^15^. In Great Britain, oak woodlands are currently threatened by Acute Oak Decline (AOD), a complex decline disease resulting from the interaction between multiple biotic and abiotic inciting factors ^16^, with similar declines of oak increasingly being reported across continental Europe, Iran and the United States ^17^. Trees exhibiting symptoms of AOD are thought to become predisposed to disease through changes in their environment: in particular, increasing drought stress is considered a key predisposing factorfor disease ^18^. Secondly, interactions with biotic agents tip the tree into a disease state. AOD is associated with the presence of larval galleries of the bark boring beetle *Agrilus biguttatus* within the inner bark ^19^, as well as inner bark tissue necrosis by a multispecies consortium of bacteria, typically consisting of *Brenneria goodwinii*, *Gibbsiella quercinecans* and *Rahnella victoriana* ^20^. These interacting factors result in a distinct decline phenotype, with thinning crowns, weeping fissures and bleeding patches rising vertically up the stem among the most conspicuous symptoms ^21^. Although the disease is widespread, not all trees at a site are affected, and trees often enter remission ^22^. This raises the question of microbial involvement in disease resistance, and particularly whether there are microbial taxa that are either indirectly improving the health of affected trees, or directly suppressing the necrotising bacteria.

Decline diseases such as AOD are conceptualised by the disease (or health) triangle ^23^, which posits that health outcomes in plants result from the complex interrelations between host factors, their microbiome (including pathogens), and the environmental conditions they experience. While the composition of plant-associated microbiomes and their importance in plant health, their interactions with plant pathogens, and their responses to abiotic variables, have been established for model plant species and crops, this has not been studied extensively for tree species that are so long-lived, play vital roles in global landscapes and are increasingly threatened by climate stressors. Environmental factors can therefore have major effects on tree health, as in AOD: changing environments can introduce stresses such as drought or mechanical damage that can weaken plants, providing opportunities for entry by pathogens or reducing host immune responses to pathogen activity ^24^. They may also have indirect effect on plant health through its effect on the plant microbiome, acting to filter for specific microbial taxa best able to survive or thrive, driving structural variation in microbiome composition and microbial function across the landscape ^25^. However, the tree microbiome has been much less intensively studied when compared with other major terrestrial and aquatic ecosystems of ecological significance (e.g. soils, oceans). Tree species harbour diverse and distinct microbiomes, and their internal woody tissues represent an ecologically important and critically underexplored ecosystem ^26^. Specific knowledge gaps include the comparative composition and function of tree microbiomes across above-and below-ground tissues, and the impact of environmental factors (e.g. rainfall, temperature, pollution) and disease faced by global tree species on microbiome composition and function. While our fundamental understanding of tree microbiome assembly, dynamics and function has been hindered by challenges associated with microbiome analyses of large, long-lived, woody tree species, comprehensive tree microbiome studies are essential to drive new discoveries and mechanistic understandings of tree-microbiota-environment interactions, and underpin translational studies to boost tree resilience to environmental change and disease.

Here, we conducted a landscape-scale microbiome analysis of 900 oak leaf (leaf punch discs), stem tissue (sapwood and phloem obtained from stem cores) and root/rhizosphere soil samples (fine roots with attached rhizosphere soil obtained from soil cores), from 300 trees (150 asymptomatic, 150 AOD symptomatic) at 30 oak woodlands across Britain (Figure 1). Using a combination of 16S rRNA gene and ITS community profiling and metagenome sequencing, we describe the taxonomic and functional composition of the above- and below-ground oak microbiome and the impact of landscape-scale environmental gradients and disease on microbiome composition and function. A three-level occupancy model was developed to detect the presence of microbial taxa at the site and tree level, enabling identification of microbiota associated with health. We also generated a metagenome assembled genome resource of the oak microbiota comprising 1657 high quality bacterial, archaeal and fungal genomes. Collectively this represents the largest landscape-scale microbiome dataset for any tree species to date, providing fundamental insights into tree microbiome composition and function across above-and below-ground tree tissues, and the impact of environmental stressors and disease on tree microbiome dynamics at the landscape scale.

**Figure 1:**
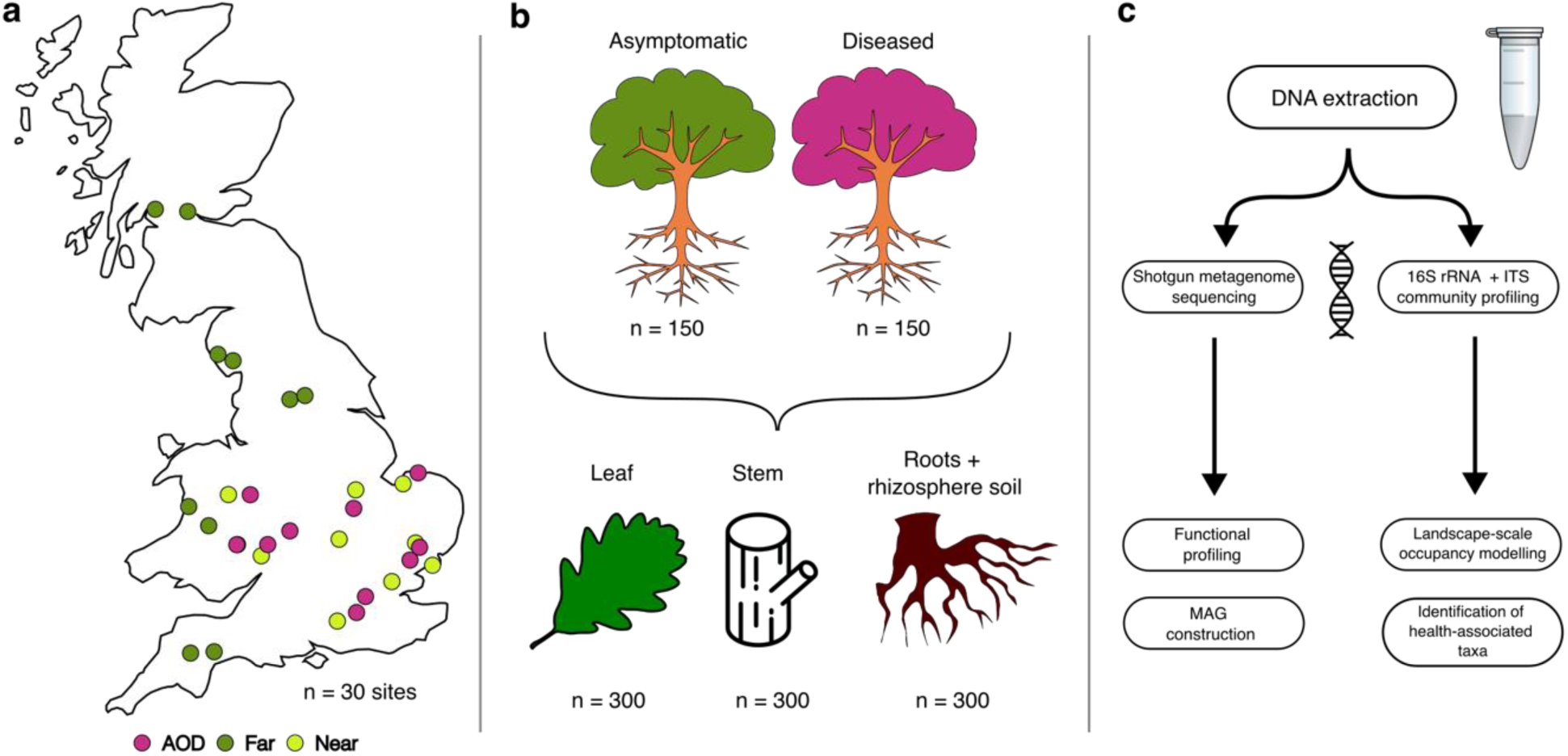
Study overview. a) Location of all 30 sampled sites across the UK. Dark green = sites outside the AOD zone, Light green = sites within the AOD zone but with no AOD symptoms present at time of sampling, Purple = sites with AOD. b) Overall sampling strategy across sites for tree disease status and tissues. c) Workflow for sample processing and analyses for each generated data set.

## Results

### The oak microbiota

Single gene community profiling of bacterial/archaeal (16S rRNA gene) and fungal (ITS) communities in 900 oak tissue samples from 300 trees across Great Britain identified 33,449 16S rRNA gene amplicon sequencing variants (ASVs) and 14,918 ITS ASVs. After filtering the dataset using read abundance thresholds, 7,806 ASVs were retained (Figure 2). Of these, 4,258 ASVs were fungal, with most ASVs belonging to orders within *Ascomycota* (2,566, 60%), and *Basdiomycota* (1,242, 29%). We identified 3,548 bacterial ASVs, across all tissue types. The majority of bacterial ASVs were identified as *Pseudomonadota* (1212, 34%), *Actinomycetota* (688, 19%), and *Acidobacteriota* (510, 14%). From the metagenome data, we recovered 15,672 mbins over 200kb in length after binning the metagenomes using VAMB ^27^, of which 1,657 were deemed high quality (a BUSCO completeness score over 50% and contamination under 10%). Of these, 180 were identified by CAT as Eukaryota, 14 as Archaea, and 1,463 as Bacteria (Figure S1). The number of bins recovered varied between tissues: most bins (1,345) were from root/rhizosphere samples (fine oak roots with attached rhizosphere soil), 119 were from stem (phloem/sapwood), and 193 were from leaves. We recovered more eukaryotic bins (126) than bacterial (67) in leaves, reflecting the higher abundance of fungal reads in leaf tissue. We obtained several genomes with particular relevance to plant health, including those of known oak pathogens, such as 9 high-quality *Brenneria goodwinii* genomes, 1 *Rahnella victoriana* genome and 1 *Gibbsiella quercinecans* genome (all associated with AOD stem canker), as well as 20 genomes of *Taphrina deformans*, a fungal foliar pathogen.

**Figure 2:**
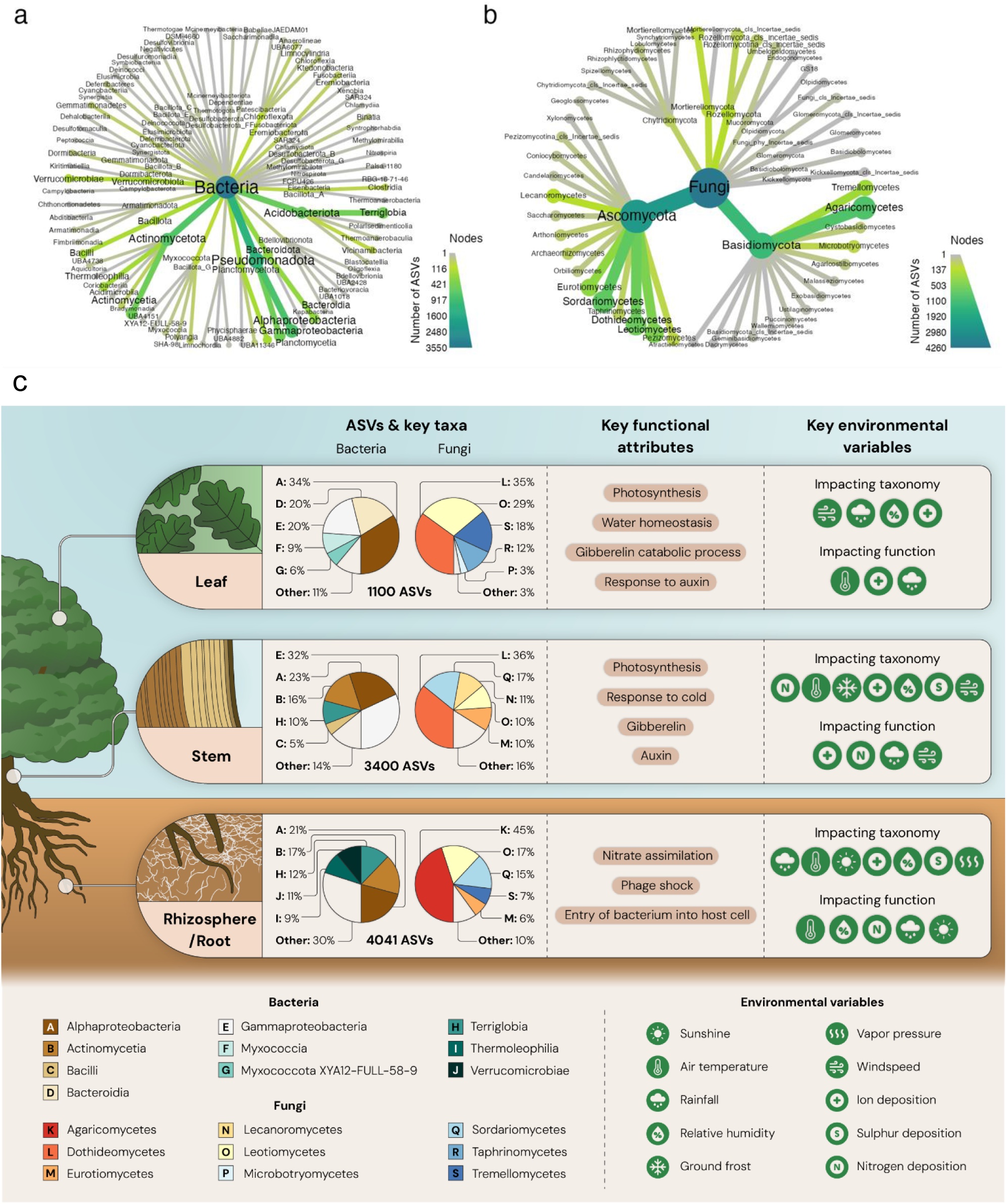
Heat trees to order level, showing the numbers of ASVs from the amplicon sequencing study for bacteria (a) and fungi (b) identified in the study across all tissue types. Diagram summarising key findings from a landscape-scale analysis of the oak microbiome (c). ASVs = ‘Amplicon Sequence Variants’ based on single gene community profiling analysis of the 16S rRNA gene (bacteria) and ITS (fungi). Key functional attributes were identified using GO term abundance in metagenome datasets. Key environmental variables impacting taxonomic and functional composition of the microbiome were identified using a three-level occupancy model.

### Tree tissues harbour taxonomically and functionally distinct microbiota

Microbial community composition differed strongly between oak leaf, stem and root/rhizosphere tissues. Leaves were much less diverse than stem and root/rhizosphere, with only 1,110 ASVs compared to 3,400 and 4,041 ASVs, respectively. The majority of ASVs were unique to each tissue type (Fig. 3C), and there were more shared ASVs among adjacent tissues on the plant (leaf and stem: 274, stem and root/rhizosphere: 347) than among non-adjacent tissues (leaf and root/rhizosphere: 20). Only 53 ASVs were present in all three tissues. Similarly, leaf samples were much more similar in their composition, with a much tighter clustering of leaf samples compared to stem and root/rhizosphere samples (Figure 3b). Of the 53 ASVs observed across all three tissue types, 46 could be assigned to genus level and belonged to 37 fungal and bacterial genera; 41% of these ASVs (n=19/46 ASVs), belonging to 11 of the 37 bacterial and fungal genera (30%) detected across all tissue types have previously been identified as oak seed endophytes that are inherited by vertical transmission ^28^.

**Figure 3:**
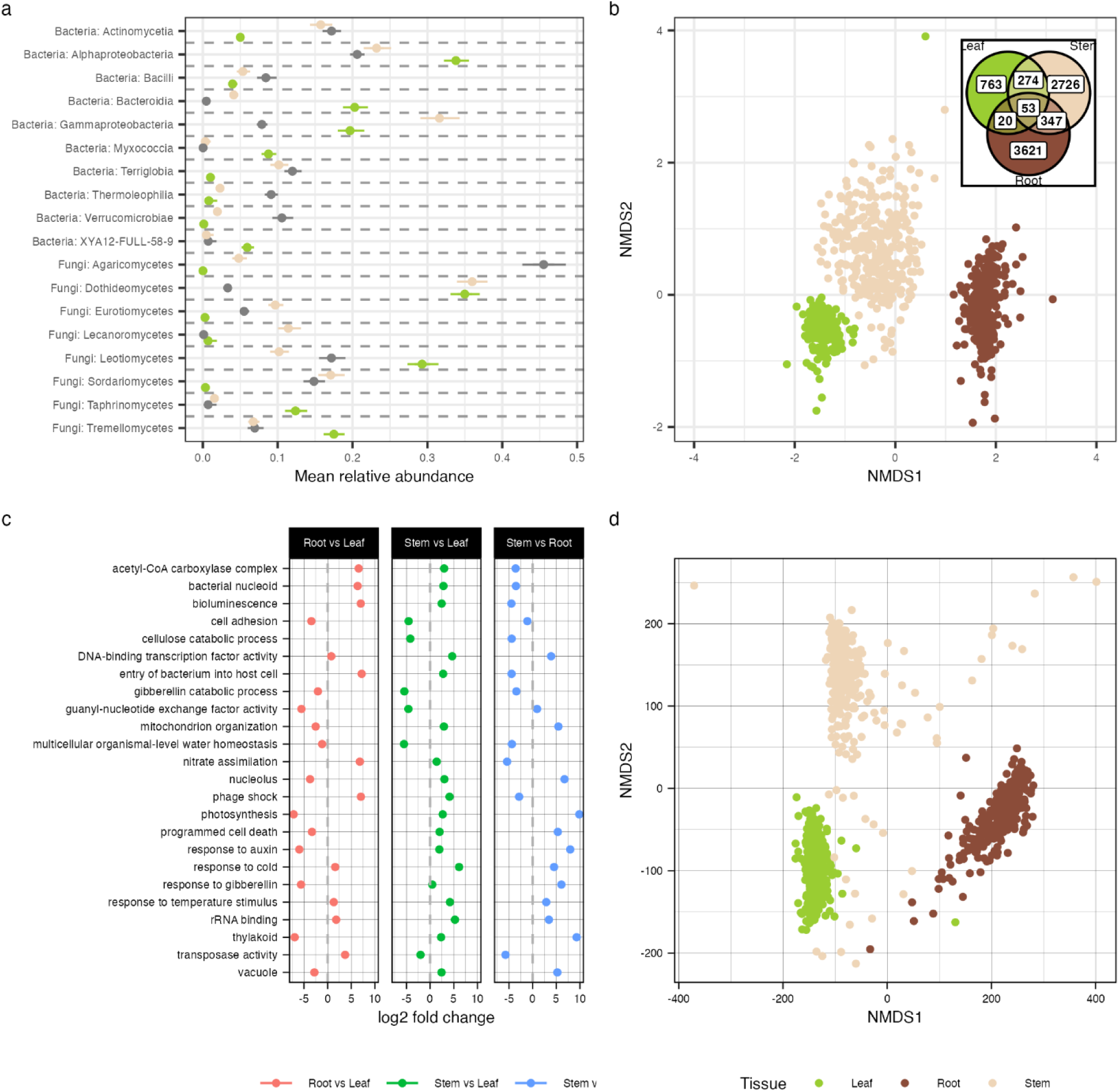
Comparison of taxonomic and functional profiles among tissue types. a) Relative abundances of taxa, aggregated at the class level, estimated using a Bayesian beta-binomial mode. Results are shown for classes where there is a significant difference (95% credible interval is not zero) in the mean relative abundance between at least one pair of tissues. b) NMDS ordination using Horn distance of the combined 16S rRNA and ITS amplicon sequencing profiles for all 900 samples. c) Highlights the largest differences in GO term abundance between each tissue, estimated in DESeq2. d) NMDS ordination using the Poisson distance for the functional profiling dataset.

At the whole community level, analysis of the filtered metagenome reads showed that leaves had a higher mean proportion of fungal reads (median RA: 0.75, 0.73 – 0.76 95% CI) compared to bacterial reads (median relative abundance (RA): 0.25, 0.24 - 0.27 95% CI) (Figure S2, Table S2). However, in all other tissues, there were more bacterial reads than fungal reads on average, particularly in root/rhizosphere, where practically all reads on average were bacterial (median RA: 0.989, 0.987 – 0.990 95% CI). To get a better breakdown of lower taxonomic levels within tissue type, we modelled the relative abundance data from the amplicon sequencing dataset (Figure 3a) using a Bayesian beta-binomial model. Leaf samples were characterised by higher mean relative abundances of Alphaproteobacteria, Bacteroidia, Myxococcia, Leotiomycetes, Taphrinomycetes, and Tremellomycetes compared to other tissues. Stem samples were characterised by high abundances of Gammaproteobacteria, Eurotiomycetes, and Lecanoromycetes. Finally, root/rhizosphere samples were dominated by Agaricomycetes, as well as Bacilli, Terriglobia, and Thermoleophilia (see Table S3 for all statistics).

These taxonomic shifts among tissues were also accompanied by strong changes in functional profiles. Each tissue type had a distinct profile, with almost no overlap among any tissues (Figure 3d). These differences were underlined by a large number of differentially abundant GO terms, with most (956 out of 1041) between-tissue comparisons estimated with DESeq2 showing evidence of significant differences at p < 0.05 in abundance between tissue types (Figure 3c). Many of these GO terms can be explained by the difference in the abundance of fungal versus bacterial components of the metagenome, particularly those relating to bacterial cell components. However, we also found strong evidence of differences among tissues that likely relate to elements of tree physiology, such as plant hormones, where responses to auxin and gibberellin were most abundant in stem tissue (all values mean log2-fold change ± 95% CI; auxin: stem vs. leaf = 1.95 ± 0.12, p = 9e-64, stem vs. root/rhizosphere = 7.94 ± 0.12, p < 2e-308; gibberellin: stem vs. leaf = 0.45 ± 0.15, p = 3.6e-3, stem vs. root/rhizosphere = 6.08 ± 0.16, p < 2e-308). Other differing terms related to changes in the physical conditions between tissues, such as water homeostasis, which was lowest in stem (multicellular organismal-level water homeostasis: stem vs. leaf = -5.53 ± 0.24, p = 2.9e-122, stem vs. root/rhizosphere = -4.38 ± 0.24, p = 4.3e-75), temperature, which was highest in stem, (response to cold: stem vs. leaf = 6.12 ± 0.07, p < 2e-308, stem vs. root/rhizosphere = 4.53 ± 0.07, p < 2e-308), and nutrient level, such as nitrate assimilation which was highest in root/rhizosphere (root/rhizosphere vs. leaf = 6.80 ± 0.23, p = 2.1e-195, stem vs. root/rhizosphere = -5.40 ± 0.24, p = 1.9e-114).

### Environment influences oak microbiome community composition and function

We developed a three-level occupancy model to describe the distribution of microbial taxa across the landscape and the probability of finding them in a sample, while correcting for the possibility of non-detection of species. Applying this to the single gene community profiling dataset, we predicted high numbers of undetected species at each site. The average number of undetected species across sites ranged from 74 (95% CI 69 – 78) ASVs for fungi in leaves to 1052 (1020 – 1085 95% CI) for fungi in stem tissue (Figure S3). Using these models, we predicted large shifts in community composition at the site level in response to the 10 landscape-level environmental predictors (Figure 4a, statistics in Table S5). In all models, rainfall and ion deposition consistently demonstrated a wide range of species responses. For other environmental predictors, we found strong differences between tissue types and the marker gene profiled. For example, the models predicted much higher turnover of bacteria (median SD: 0.44, 0.17 – 0.77 95% CI) vs fungi (median SD: 0.13, 0.02 – 0.35 95% CI) in response to N deposition, as well as windspeed (bacterial SD: 0.20, 0.03-0.43 95% CI; fungal SD: 1.03, 0.82 1.24 95% CI) and air temperature (bacterial SD: 0.33, 0.1 – 0.59 95% CI; fungal SD: 1.16, 0.88 1.48 95% CI). Using these species-level responses, we predicted the probability of ASV presence across the studied area of the UK landscape (Figure 5). These revealed varying patterns of ASV distribution across the UK landscape: for example, taxa that have a high predicted probability of occupancy in the south east of England (Figure 5a), where rainfall is lower; cosmopolitan ASVs distributed widely across the UK (Figure 5b); and ASVs that appear to be more common in western regions where rainfall is higher (Figure 5c).

**Figure 4:**
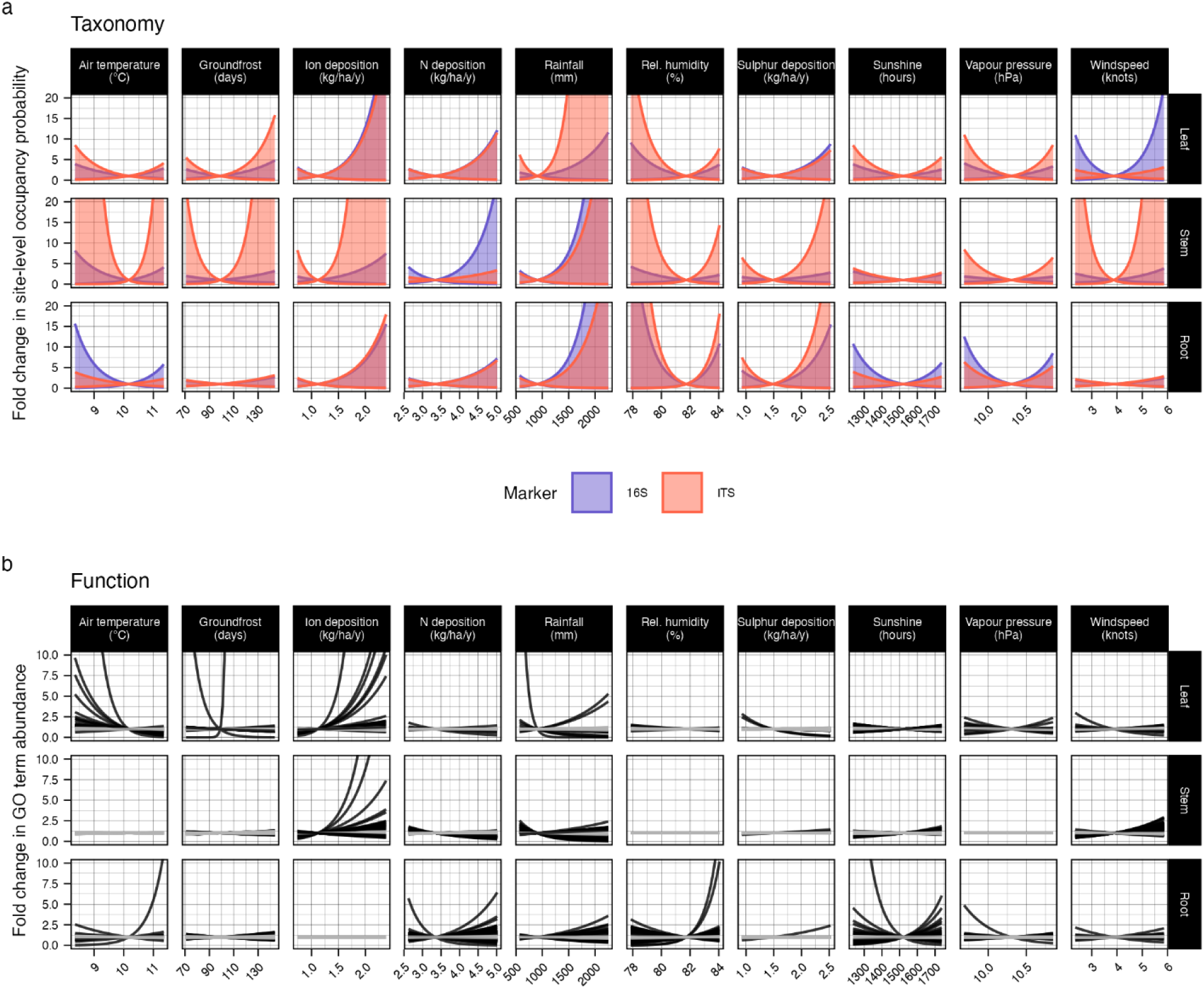
Comparison of responses of ASVs and functional terms to environmental variables within each tissue. Units for each horizontal axis are given in the column headers. a) 95% quantiles for the distribution of species responses to each environmental variable (columns) in each tissue (rows) at the site occupancy level. A wider gap between the quantiles indicates a wider range of species responses, indicating greater taxonomic shifts in response to that variable. Red intervals = fungal taxa, blue intervals = bacterial taxa. b) Response of all GO terms with a significant differential abundance (p < 0.05) to each environmental variable. Grey lines have a log2 fold change value of < 0.1, and black lines have a log2 fold change value >= 0.1.

To understand how different functional terms from the metagenomes responded to environmental variables, we modelled the abundances of functional terms using DESeq2. Of the 10,410 tissue-by-environment functional responses tested, 1,136 were significant at a p value threshold of 0.05, of which 446 had a log_2_ fold change value > 0.1. These responses, such as those for ion deposition and sunshine, often reflected the same pattern as the taxonomic responses, suggesting that shifts in taxonomic composition were related to shifts in the functions carried out by the community. Leaf tissue exhibited wide shifts in functional profiles in response to changing air temperature, ion deposition, and rainfall; stem by ion deposition, N deposition, rainfall and windspeed; and root/rhizosphere by air temperature, N deposition, rainfall, humidity, and sunshine. However, many variables had few effects which could be considered as “practically significant” (log2 fold change value < 0.1), including sulphur deposition, ion deposition in root/rhizosphere, and relative humidity in leaf and stem (Figure 4b, grey lines), often in contrast to the large range of effects in the taxonomic responses, suggesting little turnover in functional profiles for these variables, at least for the functions profiled.

**Figure 5:**
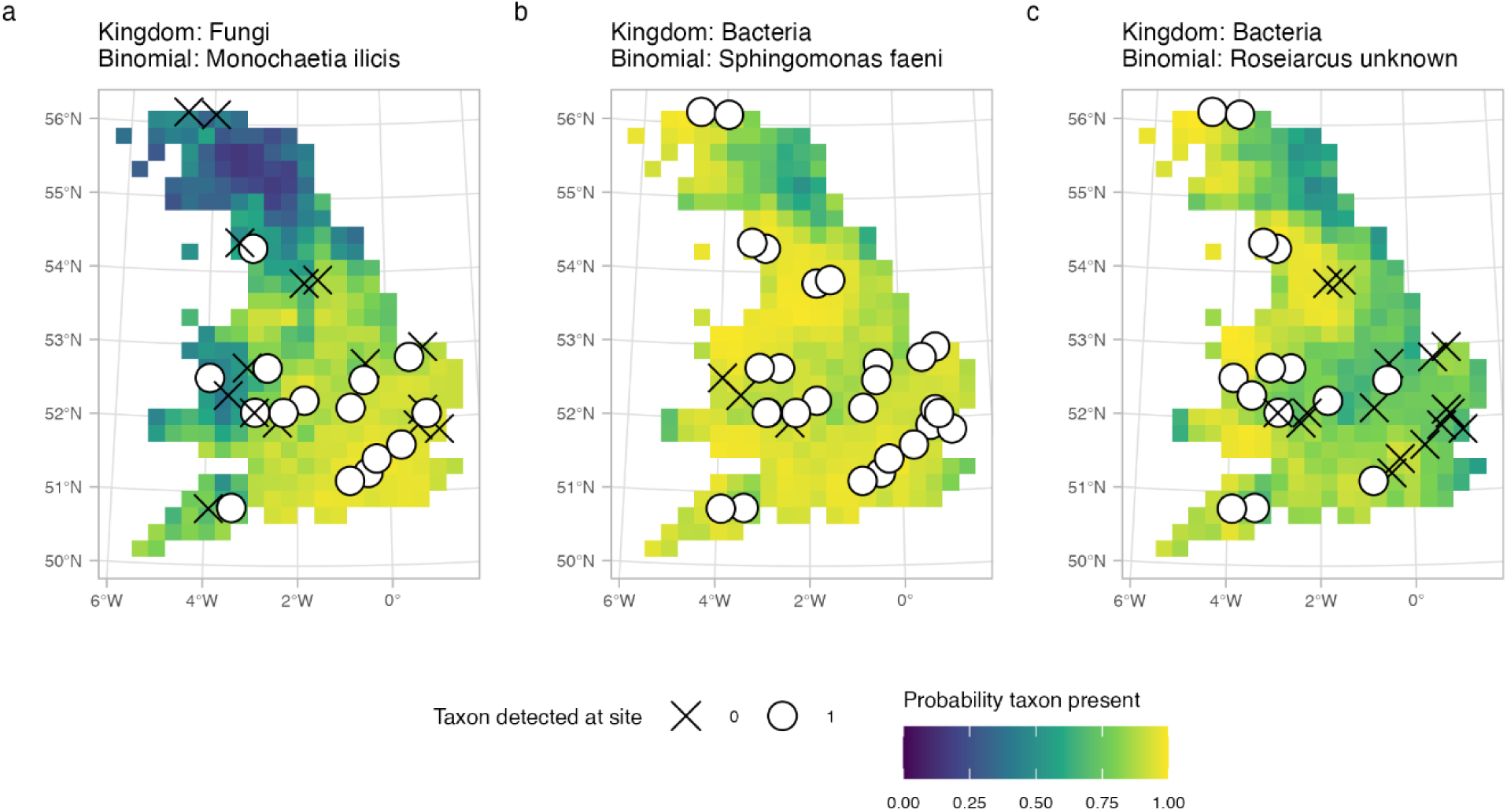
Maps showing example predicted distributions of three ASVs from a) leaves, b) stem, and c) root/rhizosphere. Map fill colour indicates the probability a site in a grid square is occupied. White circles show sites where the ASV was detected in at least 1 tree, and crosses show sites where the ASV was not found.

### Diseased stems have reduced bacterial and fungal species richness and altered microbiome function

Using occupancy models on the 16S rRNA gene and ITS community profiling data, we found little evidence of variation among species in response to disease at the sample occupancy level in leaf and root/rhizosphere tissue, but stem tissue, where bacterial canker symptoms associated with AOD are found, demonstrated a species response (Table S5). Species richness declined strongly in response to AOD in stem tissue for both fungi (-66 species, 95% CI -121 to -12) and bacteria (-105 species, 95% CI -156 to -54) (Figure 6b). At the kingdom level, metagenome analysis indicated that in stems, bacteria increased in relative abundance from 64.5% to 68.6% (median difference: 4%, 0.44 -7.7% 95% CI), with a concomitant decrease in fungal abundance. In contrast, in leaves, there was an increase in the relative abundance of fungi in diseased trees from 71.7% to 78.7% (median difference: 7.1%, 4.26% - 9.7% 95% CI) with a concomitant decrease in bacterial abundance (Table S4). For taxon abundance, we found evidence for a small amount variation in abundance of read counts in stems for bacteria (median variance: 0.41, 0.29 – 0.53 95% CI) and fungi (median variance: 0.47, 0.35 – 0.61 95% CI), and in the root/rhizosphere for fungi (median variance: 0.53, 0.43 – 0.64 95% CI)

**Figure 6:**
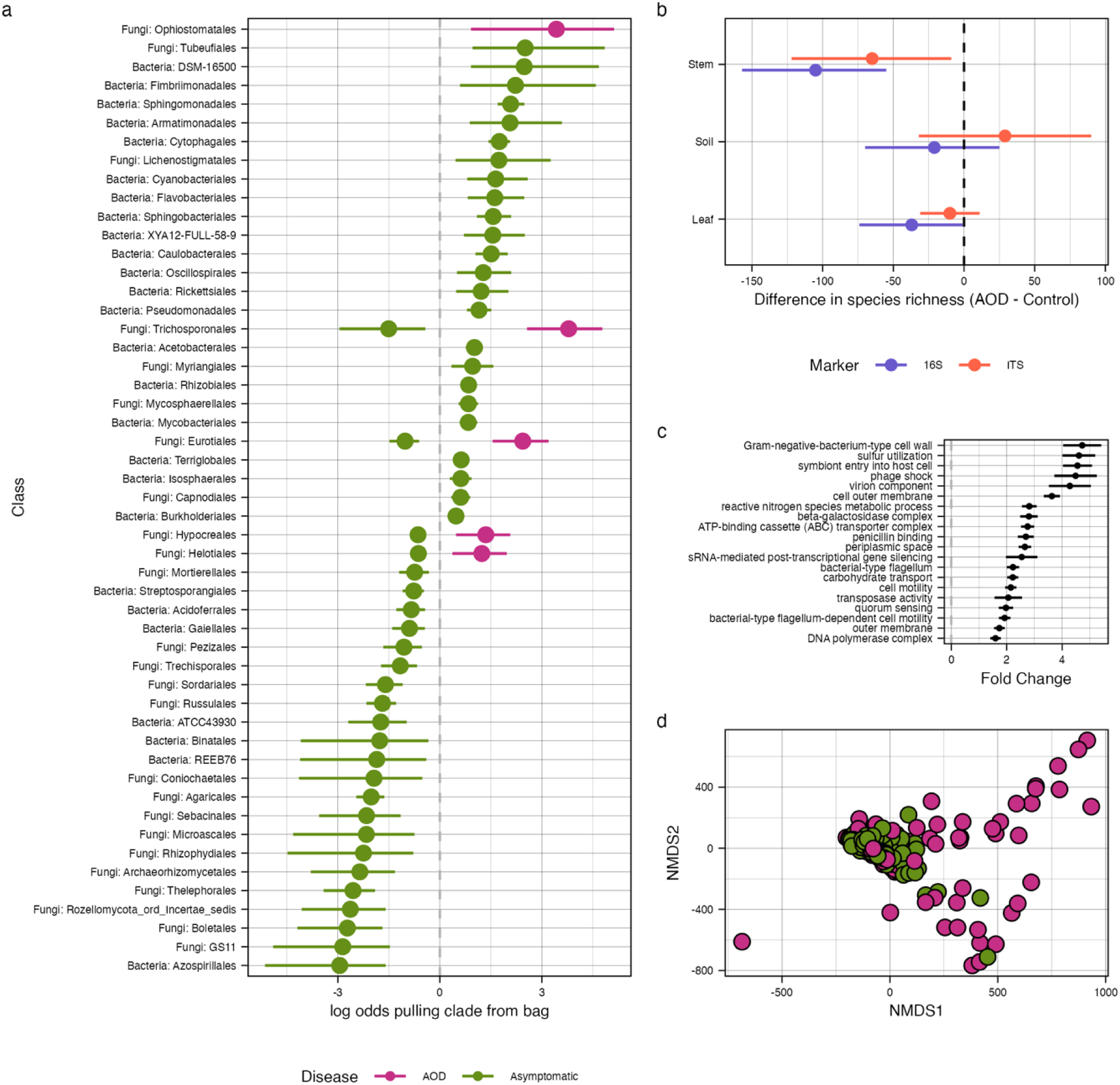
Differences in taxonomic and functional groups in response to disease. a) log-odds of finding an ASV of a given microbial class in a list of health-associated microbes (green points and lines) or diseased-associated microbes (purple points and lines). Points show mean effect size, error bars show 95% credible intervals. b) Change in estimated species richness in trees as a function of disease, estimated from the occupancy model. Here we assume a hypothetical site with average environmental values but where all species are present and vary only in their ability to occupy trees. Points show mean effect, bars show 95% credible intervals. c) Top 20 differentially abundant GO terms in diseased tree stems compared to asymptomatic tree stems, estimated with DESeq2. Points show mean effect size, error bars show 95% confidence intervals. d) NMDS ordination of stem functional profiles using the Poisson distance metric. Red points show diseased trees, and blue points show asymptomatic trees.

We identified 2,782 ASVs associated with tree health status (< 2.5% of the posterior inside a ROPE of [-0.1, 0.1]) – 34 were positively associated with AOD, and 2,748 with health (Table S6). Of the disease-associated ASVs, only one ASV was more abundant in diseased stem samples than asymptomatic stem samples (median log2 fold change: 1.71, -0.09 – 3.47 95% CI). This ASV was classified by phylogenetic placement as belonging to the genus *Brenneria*, one of the three genera of bacteria associated with AOD. All of the other disease-associated microbes were root/rhizosphere-borne fungi in the orders Trichosporonales, Ophiostomatales, Eurotiales, Hypocreales, and Hypocreales (Figure 6a). Among the ASVs associated with health, 706 were identified in leaves, 235 in root/rhizosphere, and 1,807 in stem tissue. Members of orders DSM-16500, Tubeufiales, Fimbriimonadales, Armatimonadales, Sphingomonadales, and Cytophagales were the most over-represented. Of these, Sphingomonadales were particularly abundant, with 81 of 103 ASVs positively associated with health, (median log-odds of selecting clade member: 2.08, 1.62 – 2.57 95% CI), as well as Cytophagales (92 of 127 ASVs positively associated with health; median log-odds of selecting clade member: 1.73, 1.26 – 2.15 95% CI).

All of the top 100 ASV biomarkers associated with healthy oak trees were from stem tissue, comprising 51 fungal and 49 bacterial ASVs. The top 10 health-associated ASVs with genus-level assignment belonged to *Sphaerotilus, Rhizobacter, Microcyclospora, Sphingomonas, Bradyrhizobium, Aquirickettsiella, Ramularia, Williamsia, Granulicella*. In contrast, 99 of the top 100 ASV biomarkers associated with diseased oak trees were form the root/rhizopshere and all were fungi. One ASV from the stem was a biomarker for disease, and was identified as *Brenneria goodwinii*, a key bacterial cause of stem tissue necrosis in AOD trees. The top 10 ASV biomarkers of disease with genus-level assignment belonged to *Pascua, Penicillium, Scleroderma, Fusarium, Acidomelania, Apiotrichum, Mortierella, Brenneria, Solicoccozyma, Pezicula*.

We also found that the functional profiles differed as a function of AOD status (Figure 7 - Tables S9, S10, S11). Of 1,041 tissue-disease interactions tested (highlighting differential abundance of reads between diseased and healthy trees for each GO term and tissue), 245 were significant at p < 0.05, of which 148 were in stem tissue, 56 in leaf, and 41 in root/rhizosphere. Of these, only 4 effects had an absolute log2 fold-change value > 1 (equivalent to a change in proportional abundance of 2x) in leaves, 0 in root/rhizosphere, but 38 in stem, all of which were more abundant in diseased tissue (Figure 6c). Of these, many GO terms appeared to relate to the increase in the abundance of bacteria in diseased tissues, such as Gram-negative cell walls (log2 fold-change: 4.72 ±0.7 SE, p = 3.6e-11), flagella (log2 fold-change: 2.23 ± 0.23, p = 2.13e-22), and quorum sensing (log2 fold-change: 1.97 ± 0.27, p = 3.3e-14). Other changes were related to pathogenicity, such as symbiont entry into host cells (log2 fold-change: 4.55 ± 0.54, p = 1.3e-16), as well as an increase in the abundance of virion components (log2 fold-change: 4.28 ± 0.77 SE, p = 4.9e-8). Curiously, there was also an increase in reads mapping to sulphur utilisation genes (log2 fold-change: 4.6 ± 0.6 SE, p = 1.1e-13), which has not been previously implicated in AOD. Together, 16S rRNA gene and ITS community combined with metagenomics has enabled identification of taxonomic and functional biomarkers of health and disease in oak trees.

**Figure 7:**
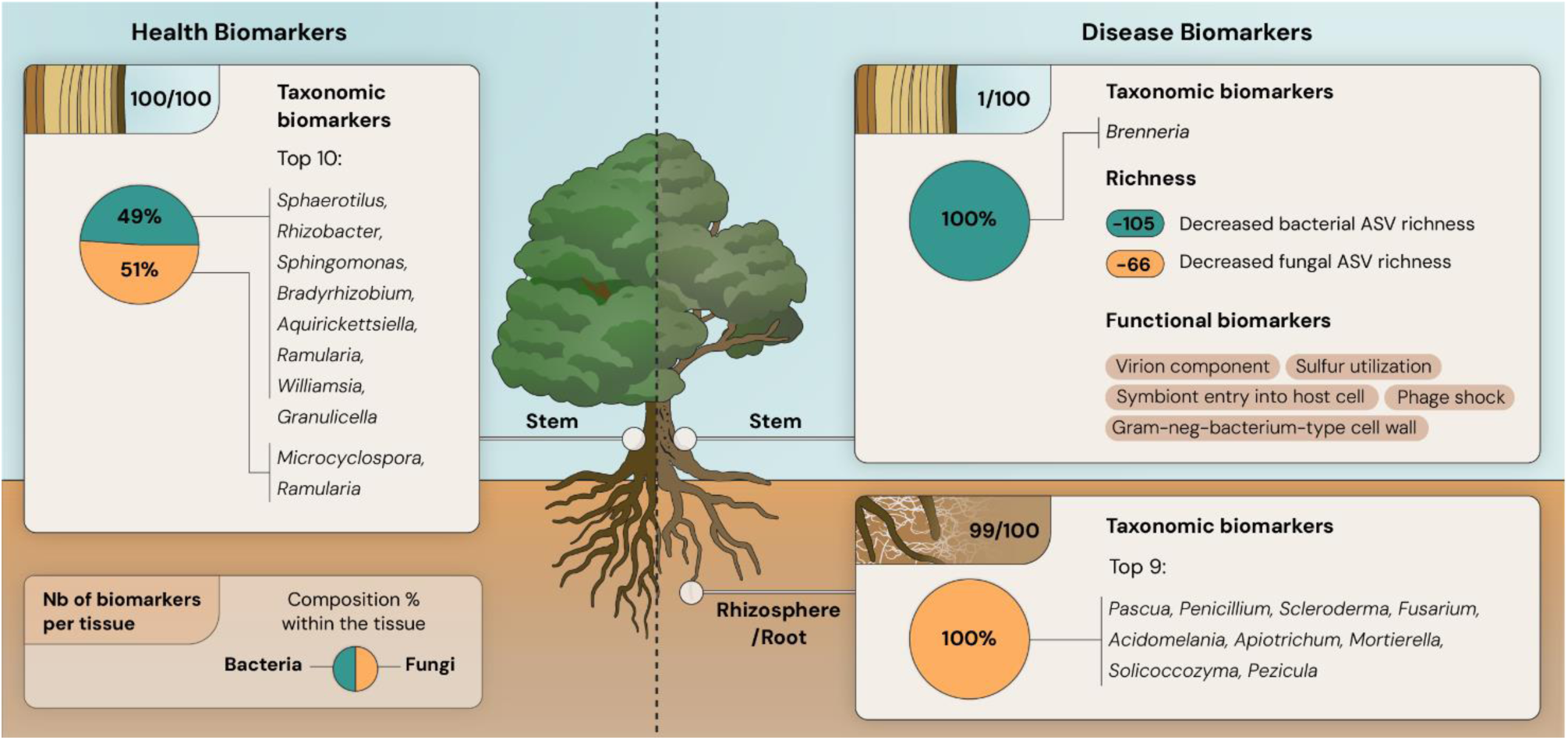
Identification of taxonomic and functional microbial biomarkers of health and disease (Acute Oak decline) in oak trees across Britain, and effects of disease on the tree microbiome. The top 100 ASVs associated with healthy and diseased oak trees were identified. The proportion of the 100 health- and disease-associated ASVs found in each tissue type is stated, along with the proportion that are bacterial or fungal, and the genus-level taxonomic assignment. The top 10 ASV biomarkers of health and disease is also shown.

## Discussion

This study represents the most comprehensive description the microbiome of a tree species to date, identifying 7,806 bacterial, archaeal and fungal ASVs, and generating 900 metagenome datasets, producing 1,657 high-quality metagenome bins, with taxonomic annotations for each species, and landscape-scale distribution models for the taxa in the amplicon sequencing dataset. Using this large dataset, we investigated the complex set of factors responsible for microbiota variation among tissue types within trees, between asymptomatic and diseased trees, and the environmental factors influencing community composition across the landscape. We also identified a list of microbial taxa most associated with plant health status, which may be candidates for experimental follow-up as biocontrol agents to treat AOD, and also represent useful targets to develop our understanding of the mechanistic processes underpinning microbiome dynamics and function oak trees.

Within trees, we found each tissue type hosted a distinct microbial community, with most ASVs unique to each tissue. In fact, there were only 53 ASVs which were common to all three tissues sampled. The number of shared ASVs between pairs of tissues was higher in tissue pairs that were closer in proximity to one another spatially in the tree: leaf and stem shared 274 ASVs, stem and root/rhizosphere shared 347, while leaf and root/rhizosphere shared only 20. Curiously, we found that compared to the other tissues, leaves hosted a higher proportion (median 75%) of reads identified as fungal. Although the direct translation from read counts to biomass is difficult due to systematic differences in genome sizes among bacteria and fungi, this may suggest that in oaks, fungi are the dominant members of the leaf microbiome. Root/rhizosphere samples contained a very small proportion of fungal reads overall (median ∼1%), reflective of prokaryotic predominance in these tissues. Compositionally, the microbial groups that were predominant in each tissue follow similar patterns to those reported in other studies, with high proportions of Proteobacteria in leaf and stem tissues, as well as high proportions of Ascomycete classes such as Leotiomycetes, Sordariomycetes, and Dothideomycetes ^26,29^.

At the landscape scale, we found that environmental variables were important predictors of community composition. Across all community types, both rainfall and ion deposition were consistently large predictors of taxon occupancy, with a wide range of plausible ASV responses indicating high levels of turnover in community composition in response to these variables. Some effects were strongly marker- and tissue-specific; for example, nitrogen deposition had its strongest effect in bacterial communities in stem tissue, and has previously been described as a predisposing factor for AOD through driving nutrient imbalances in trees and altering microbial activity ^18^. Concurrently with these changes in community composition, we found evidence for variation in functional profiles in response to the same environmental predictors. Curiously, in many of the functional tissue responses, there were few or no large effects, often in comparison to the large changes in taxonomic composition. This may indicate functional redundancy in the tree microbiome, where species are replaced with new taxa with similar function. However, it is important to note that these data came from different datasets (single gene community profiling and metagenomics), and the metagenomic data were limited in the stem and leaf samples by a large number of sequences from the oak genome.

This study provides the first analysis of variations in tree microbiome composition (comprising bacteria, Archaea and fungi) and function across above- and below-ground tissues at the landscape scale. Studies on ectomycorrhizal fungi, in oak and other tree species, have identified root/rhizosphere physico-chemical properties, air temperature, and atmospheric deposition of pollutants as key drivers of community composition ^30–32^, while a study of leaf foliar fungi in oak found evidence for latitudinal gradients in diversity and composition, though these were not decomposed to specific environmental factors ^33^. Outside of trees, a study in duckweeds (*Lemna* spp.), an aquatic plant species, identified temperature, precipitation, and nutrient concentrations as the most important environmental factors driving microbial community composition ^25^. Thus, our study provides insight into how climate and depositional variables can affect distribution of tree-associated microbiota, and extends this to cover the whole plant microbiome, rather than focusing on specific tissues.

We also found evidence for changes in the oak microbiome with disease, particularly in stem tissue, where the effects were largest. We found that species richness decreased in diseased stem tissue for both fungi and bacteria (a decrease in 66 and 105 species, respectively), concomitantly with an increase in the read count of *Brenneria* (which contains one of the causal agents of AOD, *B.* goodwinii) ASV by about 1.7x. More broadly, we found a large number (2,748) of ASVs which were more abundant in healthy trees than in diseased trees. In this species list, many microbial groups were over-represented compared to their occurrence in the whole taxon list. Of particular note were Sphingomonadales, for which 81 of 103 ASVs were found to be more abundant in healthy trees. Members of this group, particularly the genus *Sphingomonas*, have previously been reported to protect against plant pathogens in both leaves of *Arabidopsis thaliana* ^34^ and against seed-borne pathogens in rice ^35^. Members of this group may thus be good candidates for experimental follow-up as biocontrol agents against AOD. Unfortunately, we did not recover any metagenome bins belonging to this clade, which would allow investigation of the particular functional traits these taxa have.

Curiously, fungal orders containing ectomycorrhizal species, such as Russulales, Boletales, Thelephorales, and Agaricales, were all significantly under-represented in the list of health-associated microbes, indicating a robust lack of association between AOD and ectomycorrhizal fungi. Species richness and community composition of ectomycorrhizal communities has previously been shown to be unchanged between AOD and non-AOD trees ^36^. However, we found a small number of taxa that were more abundant in the root/rhizospheres of AOD trees compared to non-AOD trees. These included the pathogen *Fusarium graminearum*, various cosmopolitan root/rhizosphere-borne fungal genera including *Humicola*, *Morteriella*, and *Aspergillus*, as well as the ectomycorrhizal fungi *Scleroderma bovista* and *Tomentella ellisii*. This change in root/rhizosphere fungal communities may be related to the acidification of root/rhizospheres associated with AOD ^37^.

In addition to changes in taxonomic composition in diseased trees, we found evidence of functional shifts in the metagenome data, particularly in stem tissue, which is the primary tissue in which AOD activity occurs (Figure 6d). In diseased stem tissue, we found that 148 functional terms changed in proportional abundance. This included increases in abundance of functional terms relating to bacterial cell components, genes relating to host cell entry, and cell motility. We also found that there was a 4-fold increase in virion components, suggesting proliferation of viruses in the stem microbiome of AOD trees. Thus, the shift in function in diseased stem tissues reflects the shift to a microbiome dominated by bacterial pathogens, which in high abundance create conditions conducive to phage proliferation.

To date, there has been little descriptive work investigating the composition and function of the oak microbiome across several tissue types, and the impact of environmental factors on community dynamics. This knowledge gap limits our ability to derive mechanistic insights into tree-microbiota-environment dynamics. This work highlights the complexity of the oak microbiome and identified tissue-specific effects of environmental variables and disease on microbiome composition and function across Britain. Comprehensive tree microbiome studies are critical to understand how tree-associated microbiota respond to environmental change and disease across different tissues, to predict future climate and disease impacts on tree microbiome function, to understand their role in ecosystem function and global biogeochemical cycles, and inform translational approaches to engineer tree microbiomes for plant health.

## Methods

### Site and tree selection

We selected sites for microbiome analysis using information from previous systematic surveys of AOD across the UK. Previous work ^18^ defined an AOD ‘predisposition zone’ over south and central England using climactic variables and information on pollutant deposition to predict the probability of AOD occurrence. We selected sites in pairs: within the AOD predisposition zone, we selected sites where AOD is known to occur (AOD sites), and a nearby site with similar environmental characteristics (e.g. annual rainfall, temperature, altitude) where AOD has not previously been detected (Near sites). These sites varied in distance between each other from 1-610 km. We additionally selected 10 ‘Far’ sites lying outwith the predisposition zone for AOD, containing trees with no symptoms of AOD. For comparability, we used the same paired sampling method to select pairs of sites with similar characteristics. A map of all study sites can be found in Figure 1a.

Following site selection, we preliminarily visited all sites to confirm the presence or absence of trees with AOD symptoms, and to identify potential trees for sampling. In a small number of cases, AOD trees were found to be present at the site since the previous survey and replacement sites were found. At AOD sites, trees were classified into health status categories by counting the number of visible stem bleeds: asymptomatic trees had no visible stem bleeds on any part of the stem or branches and appeared in good physical condition, ‘low’ trees exhibited 1 to 5 stem bleeds, ‘medium’ trees had 5 to 15 bleeds, and ‘high’ trees had 16+ bleeds. At each AOD site, we selected 5 trees from each AOD category for sampling. At sites without

AOD, we sampled 5 trees that appeared in good physical condition. In total, this experimental design led to the selection of 300 trees: 150 asymptomatic oak trees and 150 AOD symptomatic oak trees (50 with ‘low’, 50 with ‘medium’, and 50 with ‘high’ symptom severity), across 30 sites in England, Scotland and Wales (Figure 1b). Selected oak trees were all mature, with a mean diameter at breast height (DBH) of 0.6 m, with a minimum of 0.2 m and a maximum of 1.5 m. A list of all sites can be found in Table S1.

### Environmental data

We extracted meteorological data for each site from the 2021 release of the HadUK dataset ^38,39^, downloading data for all 11 available variables at the 5 km grid scale. We obtained data on atmospheric pollution (total N deposition, total SOx deposition, and CaMg deposition) in forest habitats at each site using the CBED dataset ^40^ . For each variable, we assigned a value to each site using a nearest neighbour join, which assigns a site a value using the value of the nearest 5km grid cell, using the R package *sf* ^41^.

### Field sampling

Sites were visited for sample collection in the summer of 2021, from June until August. Where possible, we visited sites in order from most southernly to most northernly, to try and control for the effect of changing season, on the assumption that the southern sites reached warmer summer temperatures earlier than the more northern sites. Sampling was conducted at separate sites simultaneously by two teams, one sampling the western side of the UK and the other the east. We sampled leaf tissue from branches on the north and south sides of each tree, from the upper 25% of the crown. Using a sterile 6 mm hole punch, 3 discs were punched from each of five leaves from the north branch and south branch respectively, for a total of 30 leaf discs (approximately 0.15 - 0.2 g fresh leaf material), directly into a sterile 2ml screwcap tube. Stem tissue was sampled from asymptomatic trees, and in AOD symptomatic trees, from areas where stem tissue was asymptomatic for AOD (i.e. AOD lesion/canker tissue was avoided) using a 10 mm Osbourne Arch Punch (OAP) at approximately breast height on the north and south sides of the tree. The punched tissue was separated into its phloem and sapwood constituents, and these were both trimmed into discs of approximately 3mm depth. The sapwood and phloem discs were then cut into eighths, and one eighth from the north and one eighth from the south of each of phloem and sapwood sample were transferred to a sterile 2ml screwcap tube. Rhizosphere samples were collected using cores from a Dutch soil auger. Two cores (north and south) were taken within the drip line of each tree, and usually no more than two meters from the trunk. Root/rhizosphere cores were searched for the presence of putative oak roots, which were identified by their brown-yellow colour, and the presence of ectomycorrhizal root tips, as oaks were typically the only ectomycorrhizal host species present within range of the soil core. Oak roots (diameter 1 - 3 mm) were loosely shaken to remove bulk soil, and 250 mg of roots from each core, with adhering rhizosphere soil, were collected into a sterile 2ml screwcap tube. All equipment used for sampling was thoroughly cleaned with a 10% bleach solution between samples. All samples were immediately snap-frozen in liquid nitrogen, and then transferred to dry ice at the end of each sampling day, before being transferred to a -80 freezer.

### DNA extraction

Oak tissue samples were randomised into extraction batches within each tissue type, with each extraction batch including a negative extraction control. Extraction batches varied in size - for leaf and stem samples, batches were 29 samples, whereas for rhizosphere samples, samples were extracted in batches of 95 (with one batch of size 23 and one of size 47).

Leaf samples were homogenised by snap freezing in liquid nitrogen, and then immediately bead beating with two sterile 3 mm steel ball bearings at 2,000 m/s in a PowerLyzer 24 (MoBio Laboratories, Inc.) for 30 s. This protocol was repeated twice for each sample. Stem samples were ground with liquid nitrogen in a sterile mortar and pestle until they were reduced to small splinters. These splinters were then transferred to a 2 ml screwcap tube and homogenised using the same bead beating protocol as for leaves. DNA was extracted from this homogenised tissue as follows: 1 ml of CTAB buffer (4 % CTAB w/v, 1 % polyvinyl pyrrolidone, 0.2 M Tris-HCl (pH 8), 1.4 M NaCl, 20 mM EDTA (pH 8)) was added to each tube containing the homogenised tissue, vortexed, and heated at 60 °C for 60 minutes, shaking every 15 minutes. Samples were then centrifuged at 15,000 g for 10 minutes and the supernatant was transferred to a new tube. 4 µL of RNAse A (Qiagen 19101, 100 mg/mL) was then added, and the samples were incubated at 37 °C for 20 minutes. 800 µL of chloroform/isoamyl alcohol (24:1) was then added to each sample, and vortexed until an emulsion formed. Samples were then centrifuged at 15,000 g for 15 minutes. The aqueous phase was then separated to a new tube, and 250 µL of 6 M NaCl, 50 µL of 3 M sodium acetate, and 500 µL of ice-cold isopropanol were added to precipitate the DNA. Samples were incubated at -20 °C for 30 minutes, before being centrifuged at 15,000 g at 4 °C for 15 minutes to pellet the DNA. The pellets were cleaned with 500 µL of ice-cold 70 % ethanol, before being allowed to air dry. Once air dry, pellets were resuspended in 50 µL of TE buffer.

Rhizosphere samples were homogenised by bead beating with two 3 mm steel ball bearings, and 0.5 g of acid-washed glass beads (Sigma). Samples were snap-frozen and beaten at 2,000 m/s for 30 s, two times. 500 µL of CTAB buffer (5% w/v CTAB/phosphate buffer, 120 mM, pH 8) and 500 µL of Phenol/Chloroform/isoamyl alcohol (25:24:1) were then added to each sample. The samples were then homogenised a third time for 30 s at 2,000 m/s. Samples were then centrifuged for 15 minutes at 16,000 g for 5 minutes at 4 °C. The aqueous layer was extracted to a new tube, and 700 µL of chloroform/isoamyl alcohol (24:1) was added before vortexing to form an emulsion. The aqueous layer was again removed to a new tube, and 4 µL of RNAse A was added. Samples were incubated at 37 °C for 30 minutes. An equal volume of chloroform/isoamyl alcohol (24:1) was again added and vortexed to form an emulsion. The aqueous phase was again removed, and DNA was precipitated by the addition of 1 mL of 30% PEG 6000. Samples were then incubated at room temperature for two hours, before being centrifuged at 16,000 g for 10 minutes. DNA pellets were washed with 200 µL of ice-cold 70 % ethanol, before being air-dried and resuspended in 50 µL of TE buffer.

The resulting DNA extracts were quantified using Quant-IT Pico Green (Invitrogen, P7589) on a SpectraMax plate reader. Samples for which DNA extraction was judged to have failed (concentrations less than 0.3 ng/µL) were repeated using backup samples collected at each site.

### 16S rRNA gene and ITS amplicon PCR and sequencing

PCR was carried out separately for each tissue type and primer set. ITS primers had forward sequence ITS1F (CTTGGTCATTTAGAGGAAGTAA) and reverse sequence ITS2 (GCTGCGTTCTTCATCGATGC) ^42^, and 16S primers had forward sequence F515 (GTGBCAGCMGCCGCGGTAA) and reverse sequence R806 (GGACTACHVGGGTWTCTAAT) ^43^.

Each primer included an additional barcode at the 5’ end, with 24 available barcodes for the forward and reverse primers. This allowed us to use a combinatorial dual indexing approach, where each sample within a tissue type received a unique combination of forward and reverse barcode to allow for downstream re-identification. Both positive (*Agaricus bisporus* DNA extract for ITS; *E. coli* DNA extract for 16S rRNA gene) and negative PCR controls were run, along with all of the negative DNA extraction controls. Raw template DNA from the DNA extractions was diluted with PCR-grade water 50 times prior to PCR, as this was found to produce the greatest amplicon concentrations during preliminary testing. We used ProMega GoTaq G2 colourless master mix (catalogue number: M7833) for all PCR reactions. All PCR primers were used at a working concentration of 1 mM. For 16S rRNA PCR, we found during preliminary testing that the 16S rRNA gene primers strongly amplified oak mitochondrial and chloroplast 16S rRNA gene sequences, to proportions of approximately >99%. To combat this, we added PNA clamps specific to these two sequence types to inhibit their amplification, with sequences GTGAATTGGTTTCGAGA for mitochondria and GGCTCAACCCTGGACAG for chloroplasts, as described by ^44^.

For ITS PCR, each 25 µL reaction contained 12.5 µL 2 x master mix, 8.5 µL of PCR-quality H2O, 1 µL forward primer, 1 µL reverse primer, and 2 µL of diluted template DNA. Reactions were run on the thermocycler as follows: samples were denatured for 2 min at 95 °C, followed by then 35 cycles of the following: denaturing at 95 °C for 45 s, hold at 75 °C for 10 s, annealing at 50 °C for 60 s, and extension at 72 °C for 60 s. After cycling, reactions were held for final extension at 75 °C for 5 min, followed by a hold at 10 °C until the samples were removed from the thermocycler. For 16S rRNA gene PCR, each 25 µL reaction contained 12.5 µL master mix, 8.1 µL of PCR-quality H2O, 1 µL forward primer, 1 µL reverse primer, and 0.2 µL (5 mM) of each PNA clamp, and 2 µL of diluted template DNA. Reactions were first denatured at 94 °C for 3 min, followed by 35 cycles of: denaturation at 94 °C for 15 s, PNA clamping at 68 °C for 10 s, annealing at 50 °C for 10 s, and extension at 68 °C for 10 s. Reactions then underwent a final extension at 74 °C for 5 min, followed by indefinite hold at 10 °C.

Following PCR, all PCR amplification products were quantified using Quant-IT PicoGreen and underwent gel electrophoresis to assess the presence of PCR bands of the correct fragment length. Samples for which PCR failed were re-run, using 2 x the volume of template DNA in the reaction. We pooled samples for library prep by comparing band intensities. Bands were classified visually as ‘high’, ‘medium’, ‘low’, or ‘very low’ intensity. For each PCR plate, we then calculated the average concentration of each intensity using the PicoGreen quants, and for each plate-intensity combination, defined a volume to pool equivalent to an approximate mass of 60 ng DNA, to a maximum total volume of 10 µL. For the ITS pools, pools were cleaned up via gel extraction using the Qiagen QIAquick Gel Extraction kit (catalogue number: 28704) kit, following kit instructions. For the 16S rRNA gene pools, the same Qiagen kit was no longer available, and pools were cleaned using Agencourt AMPure XP beads according to the following protocol: 0.9x volume of beads was added to each pool and mixed by pipetting. The supernatant was then removed using a magnetic rack, and beads were washed twice using 400 µL 80 % ethanol. Beads were then dried for 5 min and resuspended in 200 µL nucleotide-free water to elute the amplicons, after which the amplicons were moved to a fresh tube. In both cases, pools were then re-combined in proportion to the number of samples they contained, to produce a single library containing roughly equal quantities of amplicons per sample. All six final libraries were then sent to Novogene for sequencing, where adapter ligation was performed, and subsequently sequenced by Illumina NovaSeq (PE250). Each library was sequenced to a depth of approximately 8Gb, with an aim of retrieving 40,000 reads per sample.

### Metagenome sequencing

For shotgun metagenomic sequencing, 30 µL of raw DNA extract for each sample was sent to Novogene for PE150 Novaseq sequencing. Leaf and stem extracts were sequenced to a depth of 25 Gb to ensure adequate coverage of the microbial fraction of the extracts, as during preliminary testing we found that 80-90 % of reads mapped to the *Q. robur* genome. Root/rhizosphere samples were sequenced to a depth of 8 Gb.

### Bioinformatics

#### Amplicon sequence data processing

The 16S rRNA gene and ITS amplicon sequencing data were first demultiplexed using Cutadapt ^45^, using an error rate of 0.15 and allowing no indels. The demultiplexed data was then processed using the nf-core/ampliseq pipeline (v2.7.0, doi: 10.3389/fmicb.2020.550420) of the nf-core collection of workflows ^46^. The pipeline was executed with Nextflow v22.10.6 ^47^. Briefly, data quality was first evaluated using FastQC (Andrews, 2010, v0.12.1). Primer sequences were then trimmed from the raw data using Cutadapt with sequences without primer sequences being considered as artefacts, with a minimum overlap of 3 and an error rate of 0.1. On average, 51.4% (+/-25.5% SD) of reads per sample passed the Cutadapt filtering stage for the 16S rRNA gene libraries, and 46.2% (+/-29.75% SD) for the ITS libraries. The sequences passing the Cutadapt filter were then processed in R with DADA2 (Callahan et al., 2016, v1.28.0), in pooled mode for each sequencing run. Reads were trimmed at the 3’ end for 16S rRNA gene libraries only (to 200 bp forward and reverse) and reads with >2 expected errors were discarded. Reads were then error-corrected, merged, and PCR chimeras were removed using DADA2, resulting in 33,449 amplicon sequencing variants (ASVs) for the 16S rRNA gene samples, and 14,918 ASVs for the ITS samples. After initial processing, for 16S rRNA samples, between 3.67% and 95.59% reads per sample (average 86.2%) were retained. ITS samples, between 4.36% and 89.47% reads per sample (average 71.5%) were retained.

Taxonomic classification of each ASV was performed for both marker genes using the DADA2 naïve Bayesian classifer; ITS amplicons were classified using the UNITE Fungi database ^50^, and 16S rRNA gene amplicons were classified using the Silva prokaryotic SSU database ^51^. 16S rRNA gene ASVs were also classified by phylogenetic placement using the nf-core/phyloplace pipeline with the SBDI-GTDB database ^52^. We used two methods as we found that the phylogenetic placement method provides more accurate classification but is unable to classify mitochondrial and plastid sequences as these are not in the reference database. We thus used the DADA2 classifications for all filtering steps, and the phylogenetic placement classifications for everything else.

We filtered the resulting data to remove non-target species: first, ASVs which were unable to be classified as either bacteria or fungi at the Kingdom level were filtered from the dataset. Second, for the 16S rRNA gene dataset, we filtered out ASVs identified as either chloroplast or mitochondria. To reduce the complexity of the dataset for downstream modelling, we then applied an abundance threshold to the ASV dataset, keeping only ASVs with at least 100 reads (ITS) or 50 reads (16S rRNA gene) within at least one sample. After filtering, the ITS dataset retained 4,258 ASVs, and the 16S rRNA gene dataset retained 3,548 ASVs. Between 0% and 98.9% of reads (average 85.4%) were retained in the ITS dataset, and between 0% and 95.1% (average 26.4%) were retained in the 16S rRNA gene dataset. The numbers of reads remaining after each filtering step during sequence processing can be seen in Figure S4.

#### Metagenome processing, functional profiling and MAG binning

Following library preparation and sequencing, we obtained 881 total shotgun metagenomes out of 900 sent for sequencing (298 leaf, 300 stem, 283 root/rhizosphere). These metagenomes were first quality-assessed and filtered to remove host (oak) DNA sequences. First, the data quality of each shotgun library was assessed using FastQC ^48^. Libraries were then processed by fastp (Chen, 2023, v0.23.2) to remove low quality reads and trim adapters. For a sequence read to be kept, a minimum of 40 % of bases were required to be above a phred score of 15. Reads were trimmed using sliding windows using the options ‘--cut_front’, ‘--cut_tail’, and ‘--cut_mean_quality 15’. Finally, reads shorter than 15 bp following this process were dropped. Next, reads mapping to the *Quercus robur* genome were filtered from each library using bowtie2 (Langmead et al., 2019, v2.4.2). For this, we used the genome from the Darwin Tree of Life project (GenBank ID: GCA_932294425.1), with bowtie2 in “sensitive” mode. Reads not aligning concordantly with the genome were kept for downstream analysis. After preprocessing, we filtered on average 85.8% of reads from leaf libraries, 87.4% of reads from stem libraries, and 0.52% of reads from root/rhizosphere libraries (Figure S5).

Because we filtered out the majority of reads from the leaf and stem tissue samples, we decided to co-assemble the libraries to pool reads from similar sources and improve overall assembly quality. We thus pooled libraries by tissue and site for co-assembly, resulting in 90 total assembles. Libraries were assembled using MEGAHIT (Li et al., 2016, v1.2.9). We then taxonomically classified each contig in each assembly using the MMSeqs2 taxonomy workflow (Mirdita et al., 2021, v14.7e284) using the UniRef90 database, which revealed that despite filtering reads that mapped to the host genome, there were still many contigs that were identifiable as either *Q. robur* or more generally from plants. To remove contigs originating from the oak genome from further analysis, we filtered the resulting database to remove any contig classified as belonging to the clade Viridiplantae (NCBI Taxon code: 33090). Statistics for the filtered assemblies are available in Table S7.

To functionally profile each metagenome, we first predicted the genes in each metagenome assembly with MetaEuk (Karin et al., 2020, v6.a5d39d9), using its “easy-predict” workflow with the eggnog database ^58^. We then clustered the gene predictions from all assemblies for each tissue using the MMSeqs2 easy-cluster workflow, with a minimum sequence identity of 95% and an alignment coverage of query to target of 90%. We then annotated the centroid of each cluster using eggnog-mapper (Cantalapiedra et al., 2021, v2.1.12-0) in MMseqs2 mode, and counted the number of reads aligning to each gene prediction using CoverM (Aroney et al., 2024, v0.7.0). Finally, we constructed a list of Gene Ontology (GO) terms ^61,62^ of interest by combining the terms from three GO slims ^63^: metagenome, prokaryote, and yeast. We added an additional set of manually selected GO terms to this list that we thought might relate to the collected environmental variables, such as drought tolerance and nitrogen use. For each selected GO term for each library, we summed the number of reads mapping to genes annotated with that GO term, to estimate the abundance in each sample of each functional annotation. We also counted the number of reads mapping to genes with no GO terms (“unannotated”), and the number of reads not mapping to any genes (“unmapped”).

To reconstruct the individual genomes comprising each metagenome, we binned contigs with VAMB (Nissen et al., 2021, v4.1.4.dev105+g24bab51), using the recommended “multi-split” approach, where metagenome assemblies are concatenated together, but bins are split from samples individually. For binning, we discarded all contigs under 2000 bp in length, and mapped the combined reads from each site to the combined assembly using minimap2 (Li, 2021, v2.28), before running VAMB with default parameters. We assessed the quality of the output bins over 200 kb in length using BUSCO (Manni et al., 2021, v5.4.3) in auto-lineage mode, marking bins with a completeness score over 50% and contamination score under 10% as “high quality”. Where available, we used the specific lineage score over the domain-level score for this. We taxonomically classified all high-quality bins using BAT from the CAT_pack package (von Meijenfeldt et al., 2019, v6.0.1) using the supplied nr database (dated 2024/04/29), and additionally classified all bins nominally identified as bacteria by BUSCO using GTDB-Tk (Chaumeil et al., 2022, v2.4.0) with the database release R220. Data on all bins is available in Table S8.

### Statistical analysis

All statistical analyses were carried out in R 4.3.2 ^69^. ASV and sample data was imported into R using phyloseq (McMurdie and Holmes, 2013, v1.48.0). All Bayesian analyses were carried out using Stan (Stan Development Team, 2024, v2.34.1), either via cmdstanr (Gabry et al., 2024, v0.8.1) or brms (Bürkner, 2017, v2.21.0). We used the posterior package to post-process draws (Bürkner et al., 2023, v1.5.0).

#### Tag jumping correction

Upon inspection of the amplicon sequencing data, it became clear that tag jumping ^75^ was an issue during library preparation, which due to the combinatorial dual indexing used in the study, allowed sequences to jump between samples, leading to false-positive detections. To correct the data for these false positives, we modified the approach from Rodriguez-Martinez et al., (2023). Assuming the data from unused tag jumping combinations represented an unbiased measurement of tag jumping rates, we fit a Bayesian negative binomial regression model for each sequencing library in brms to model the expected rate of tag jumping in these unused tag combinations for each ASV. We captured the geometric element of tag jumping by considering the tag matrix, the n x n matrix of the read counts for each pairwise tag combination. We modelled the tag jumping rate as the log of the number of reads in the tag matrix column plus the log of the number of reads in the tag matrix row. The model was hierarchical, with a varying intercept and slope for each ASV. We used weakly-informative priors on all parameters to constrain the model, with a normal(0, 3) prior on the fixed effects, an exponential(2) prior on the varying effect standard deviations, and an lkj(3) prior ^77^ on the correlation between the varying intercept and slope. For each used tag combination and ASV, we then predicted the upper 95% CI for the number of reads expected if there were no reads present originally in that combination. If the observed number of reads was higher than this limit, we kept the data, and if it was lower, the reads for that ASV in that sample were set to 0. This allowed us to take a conservative approach, retaining only counts where true positives were highly plausible.

#### Landscape-scale occupancy model

To analyse the data arising from the landscape-scale amplicon sequencing survey data while accounting for the possibility of imperfect detection of microbial taxa, we adapted the three-level occupancy model approach of Fukaya et al., (2022). This approach recapitulates the hierarchy of samples in our study, considering occupancy at the site and then individual tree level, and using sequencing reads from individual samples as a detection model. First, we model the occupancy of taxon *i* at site *j* (z_ij_) with probability ѱ_ij_ which we allowed to vary as a function of environmental predictors:

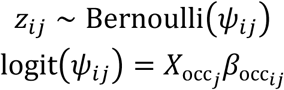

Next, we consider the occupancy by taxon *i* of sample k within site *j* (u_ijk_) with probability θ_ijk_:

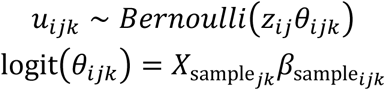

Finally, we consider the abundance of sequencing reads belonging to taxon *i* within sample *k* at site *j*. Here we depart from the approach from Fukaya et al. (2022) and model the number of reads *Y_ijk_* with a negative binomial rather than multinomial distribution, which allows easier marginalisation of the latent variables z_ij_ and u_ijk_. For each sample, we include a sample-specific size factor *a_jk_* to account for variation in sequencing depth among samples, and a taxon specific overdispersion parameter *ϕ_i_*:

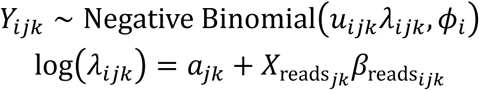

At all levels, we included a taxon-specific intercept to allow taxa to vary in their overall abundance. We additionally included taxon-specific responses to the available environmental variables at each level, and taxon-specific responses to disease status at the sample occupancy and abundance levels. We allowed all taxon-specific components to vary hierarchically, to allow pooling of information among ASVs. For each variable, we modelled a single hierarchical variance parameter, along with a simplex variable that described the proportion of the total variance attributable to each level of the sample hierarchy. We also included a single correlation matrix to model correlations among the environmental variables, assuming that the correlations among variables should be the same across levels.

We applied weakly informative priors to all model components in order to constrain the parameter space to values within the realm of biological plausibility. For standard deviation parameters, we chose exponential(2) priors, and for the simplex variables we used uniform dirichlet(1, 1, 1) priors for the environmental variables and a beta(1, 1) prior for the disease variable. For the means of the hierarchical distributions, we used normal(0, 0.5) priors for occupancy means and normal(0, 1) priors for the abundance means. This constrained taxon-specific proportional odds shifts to values between 0.17 and 4.8x, and proportional abundance magnitudes between 0.1 and 11x per unit. For the correlation matrix, we applied an LKJ(3) prior, which provides weak regularisation to keep plausible correlations among species parameters between -0.48 to 0.48 (95% CI). Finally, we placed a lognormal(0, 1) prior on the size factors *a_jk_* to ensure they were strictly positive, enabling their interpretation as an offset.

This model was implemented in Stan (version 2.34.1) and fit using the cmdstanr package. All input environmental variables were scaled to have a mean of zero and unit standard deviation, aiding model convergence and allowing easier prior specification. Due to restrictions in computational resources, we fit one model for each combination of marker gene and tissue type, for 6 total models. Using the posterior distribution, we predicted the true occupancy of each taxon at each site and sample, conditional on the detection history of that sample unit, and subsequently the true species richness of each sample unit. We also predicted the distribution of each ASV across the UK landscape using a 25 km gridded dataset covering the whole of the UK.

Using the log-fold changes in read abundances between asymptomatic and diseased trees, we defined ‘health-associated’ microbes as those for which the <2.5% of the posterior distribution was within the region of (-0.1, 0.1). These were split into two groups, health- and disease-associated, by the sign of the median of the posterior. To examine over-representation of specific taxonomic orders in this list, we built a Bayesian extended hypergeometric model to estimate the log odds ratio of selecting a member of a given order, given the number of taxa of that order in the whole dataset ^79^.

#### Functional profiling of metagenomes

To visualise differences among tissues in their functional profiles, we first conducted an NMDS ordination of all samples using the metaMDS function in the vegan package ^80^, using the Poisson distance metric implemented in the PoiClaClu package ^81^. To model the abundances of reads mapping to genes annotated with specific GO terms, we used DESeq2 ^82^ to model log-fold changes in abundance as a function of terms of interest. To model log fold-changes among tissue types, we constructed a model with an interaction between tissue type and tree disease status; this allowed us to estimate log2 fold-changes in abundance only in asymptomatic trees. To compare functional profiles as a function of disease, we constructed per-tissue models featuring tree disease status, as well as all 10 environmental variables. Finally, to explore the relationship between functional terms and the environment, we constructed per-tissue models containing each of the environmental variables. To correct for multiple comparisons, we applied a false discovery rate (BH) with a nominal significance threshold of 0.05; where we ran per-tissue models, we applied this correction over the terms from all models simultaneously.

#### Tissue comparison

To compare the microbial communities of each tissue, we first partitioned the ASV table into numbers of ASVs unique to each tissue type and the numbers of ASVs shared between any two tissues, or all tissues, for each marker gene. To visualise broad difference in community composition, we conducted an NMDS ordination on the combined data for both marker genes, using the phyloseq ordinate function with the Horn index, as this index is not susceptible to variation in sampling depth ^83^. We measured the degree of association between abundance at a given taxonomic level (kingdom and class) and tissue using Bayesian beta-binomial models, with tissue-specific means and overdispersion parameters. We placed a joint multivariate-normal prior on these with a mean of zero, a variance of 0.5, and a correlation of 0.4, expressing a prior expectation that where the means relative abundance was low, there would be less overdispersion in the data. Data for proportions at the class level came from the amplicon sequencing dataset, and at the kingdom level from the metagenome reads mapped to the predicted genes from the functional profile analysis.

## Data Availability

Sequence data that support the findings of this study have been deposited in the European Nucleotide Archive (ENA) under the ENA bioproject PRJEB95962.

## Code Availability

Code for the statistical analyses, as well as the data artefacts necessary to run the analyses, are available on GitHub. The repository for the amplicon analysis is available at https://github.com/prototaxites/fo_occupancy_model, and the repository for the functional analysis is available at https://github.com/prototaxites/fo_functional_analysis.

## Acknowledgements

This work was funded by the UK Research and Innovation’s (UKRI) Strategic Priorities Fund (SPF) program on Bacterial Plant Diseases (FUTURE OAK project; grant BB/T01069X/1) funded by the Biotechnology and Biological Sciences Research Council (BBSRC), Natural Environment Research Council (NERC), Department for Environmental Food and Rural Affairs (DEFRA), and The Scottish Government. We thank the Forestry Commission’s Technical Services Unit (TSU) staff, especially Liz Richardson, Mark Oram and Alistair MacLeod for assistance with sample collection. We also thank the woodland site owners for permissions to sample oak trees at sites across Britain. We are grateful to Amy Ellison for helpful advice and discussions regarding amplicon sequencing.

## Author contributions

**Jim Downie:** Conceptualisation, Methodology, Software, Formal analysis, Investigation, Data Curation, Writing – Original Draft, Visualization, fieldwork, project management, supervision, **Alejandra Ordonez:** Investigation, Writing - Review & Editing. **Marine C. Cambon:** Investigation, Writing - Review & Editing. **Usman Hussain:** Investigation, Writing - Review & Editing. **Nathan Brown:** Conceptualisation, Resources, Writing - Review & Editing, Funding acquisition. **Manfred Beckmann:** Conceptualisation, Resources, Funding acquisition. **Jasen Finch:** Conceptualisation, Resources. **John Draper:** Conceptualisation, Writing - Review & Editing, Resources, Funding acquisition. **Sandra Denman:** Conceptualisation, Resources, Methodology, Writing - Review & Editing, Project administration, Funding acquisition. **James E. McDonald:** Conceptualisation, Methodology, Investigation, Writing – Original Draft, Visualization, Supervision, Project administration, Funding acquisition.

## Competing interests

The authors declare no competing interests.

## Notes

### Competing Interest Statement

The authors have declared no competing interest.

